# A prebiotic protocell model based on dynamic protein membranes accommodating anabolic reactions

**DOI:** 10.1101/463356

**Authors:** Andreas Schreiber, Matthias C. Huber, Stefan M. Schiller

**Author notes:** These authors contributed equally to this work (Andreas Schreiber, Matthias C. Huber). Corresponding authors: Stefan Schiller; Matthias C. Huber; Andreas Schreiber.

## Abstract

Phospholipid membranes are essential constituents of extant cells rendering them preferred candidates as membrane components in origin of life scenarios. These models greatly neglect stability requirements and their problematic synthetic complexity necessary to access such lipid membrane constituents under early life conditions. Here we present an alternative protocell model, based on amphiphilic protein membranes constituted of prebiotic amino acids. These self-assembled dynamic Protein Membrane Based Compartments (PMBC) are impressively stable and compatible with prevalent protocell membrane constituents. PMBCs can enclose functional proteins, undergo membrane fusion, phase separate, accommodate anabolic ligation reactions and DNA encoded synthesis of their own membrane constituents. Our findings suggest that prebiotic PMBC represent a new type of protocell as plausible ancestor of current lipid-based cells. They can be used to design simple artificial cells important for the study of structural and catalytic pathways related to the emergence of life.

## Introduction

The study of the emergence of life and underlying fundamental principles such as compartmentalisation and self-replication attracts great attention since decades. Self-assembly, compartmentalisation and replication are essential for the evolution of self-sustaining protocells. The prebiotic protocell requires three functional molecular species: information-storing molecules capable of replication such as RNA, catalysts e.g. ribozymes structurally encoded by that information, and compartment-forming molecules. The information replication requires at least two RNA molecules - a replicase (ribozyme) acting as the polymerase and a second RNA molecule acting as a template^1^. To keep replicase and template in proximity some form of compartmentation is essential. In addition, replication on the compartment level is required for the selection and evolution of more efficient compartments and replicases. Therefore the boundary molecules must potentially be replicable and most notably be easy to synthesize. These requirements are meet by simple peptide repeats much better than by lipids. The first reaction compartments and protocells are widely assumed to be surrounded by amphiphilic lipid-based membranes^2,1,3,4^, since most cellular compartments and cells we know today are based on lipid membranes. However, there are crucial limitations of lipid bilayer membranes of protocell models: Phospholipid membranes are highly effective barriers to polar and charged molecules, necessitating complex protein channels and pumps to allow the exchange of these molecules with the external environment. Contemporary phospholipid membranes are non-permeable to essential molecules for life, growth, and replication^5^. Second, lipid bilayer membranes lack the dynamic properties required for both membrane growth and the intake of nutrients^6^. Third and most important, so far there is no plausible way of lipid synthesis^7,8,9^ described under prebiotic condition in recent literature. Thus, even though lipid vesicles are prominent models in synthetic protocell research^10,11,12,13,14,15^, the conceptual link between the RNA replicase, and compartmentalization strategies based on lipid bilayer membranes is still lacking. Alternative models of prebiotic compartments are based on fatty-acid vesicles^16,5,6^, protein microspheres^17^, nanofibers^18^ and membrane-free compartments such as peptide-nucleotide microdroplets^19^ and polyamine - DNA aggregates^20^. Here we propose a prebiotic PMBC protocell model based on peptide repeats constituted of prebiotic amino acids (aa). Simple amphiphilic block domain proteins can be created based on pentapeptide repeats consisting of only four different prebiotic amino acids for each repeating unit such as (VPGHG)_20_(VPGIG)_30_ assembling to PMBCs *in vitro*. These extremely stable protocell models withstand temperatures up to 100°C, pH 2-11, 5 M salt and at least 8 month of storage at room temperature. They are compatible with current protocell model membrane constituents such as fatty acids, lipid precursors^6^ and contemporary phospholipid-based membranes. These cell-sized PMBCs can obtain cellular functions by encapsulating small molecules up to intact proteins and complex anabolic reactions. The exchange of vesicular content can be observed by dynamic membrane fusion. These PMBC protocell models reside anabolic ligation reactions of short DNA with 15 bp up to 1200 bp oligomers. In addition and in contrast to the complex phospholipid synthesis, PMBCs embed the synthesis of their own membrane constituents through *in vitro* transcription and translation and incorporate this amphiphilic protein building block into the membrane. The emergence of PMBCs and their self-assembly as possible primary prebiotic protocells and associated catalytic processes are illustrated in Figure 1.

**Figure 1:**
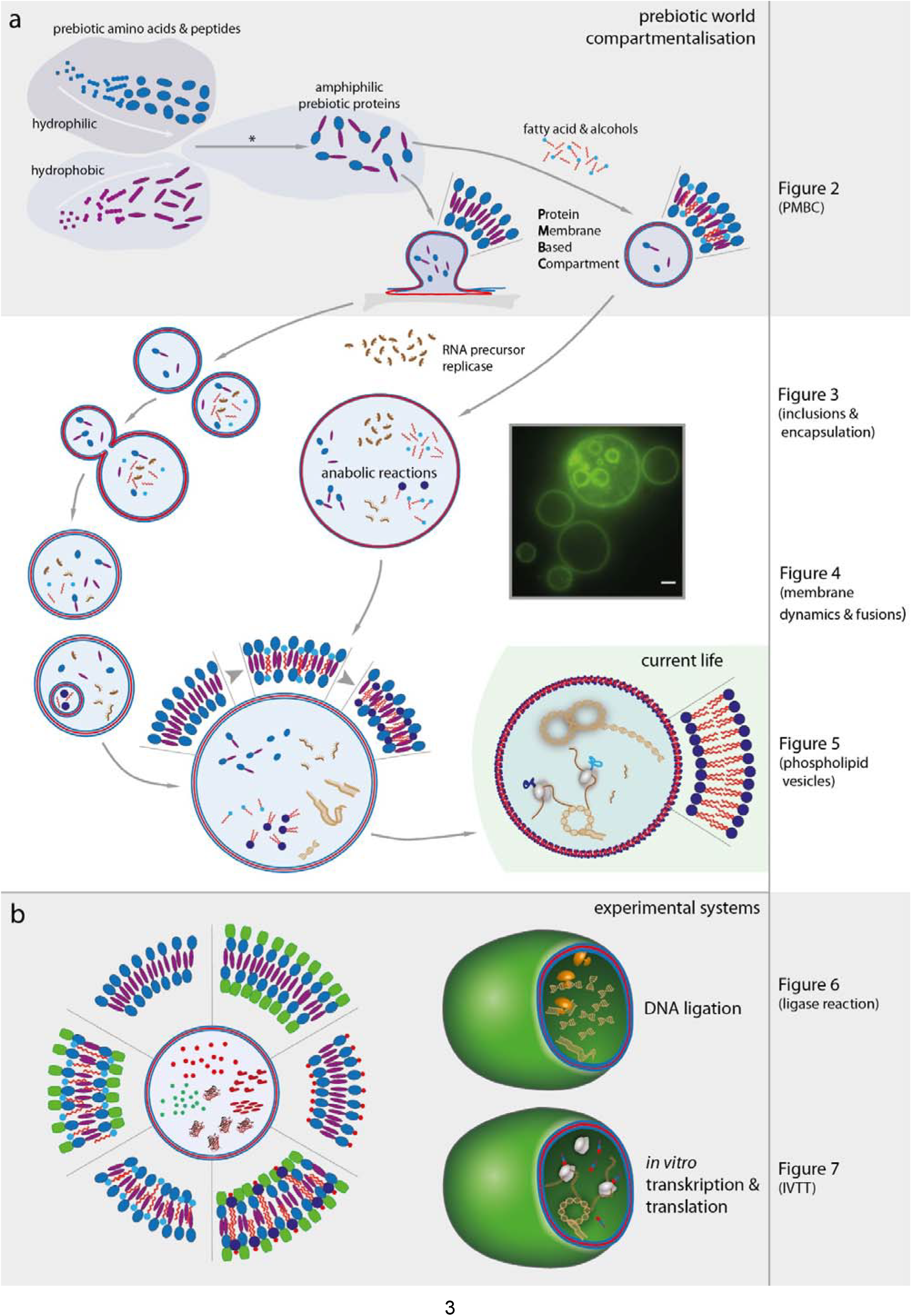
PMBCs and their self-assembly as possible prime prebiotic protocells and associated catalytic processes. **a**, **grey panel** Illustration of the chain elongation and amphiphilic peptide formation based on prebiotic hydrophobic (violet symbols) and hydrophilic (blue symbols) aa^21^. Oligopeptides in the prebiotic era have been shown to be synthesized on different inorganic substrates ^22,23^, as well as through their catalytic^24^ or auto-catalytic^25^ activity (Fig. 1a). The respective peptides or (auto-) catalyzed interconnection of hydrophilic and hydrophobic proteins (Fig. 1a, asterisk) will then lead to amphiphilic protein block-copolymers that are able to self-assemble into PMBCs. Our results indicate two prebiotic PMBC assembly scenarios. One is solely based on swelling of rehydrated amphiphilic peptides in water and a second is based on a mixture of amphiphilic proteins with small fraction of fatty alcohols (see Fig. 2, right column). Fatty acids and their corresponding alcohols can be synthesized under prebiotic conditions^26,27^ and found on meteorites in high abundancies^28^ making fat alcohol triggered PMBC assembly possible. **white panel**, The left side illustrates the properties and development of PMBCs as first entity and encapsulating other components necessary on the path towards the emergence of living evolvable systems. Once, simple prebiotic amphiphilic peptides are self-assembled into PMBCs, they can enclose short peptides, inorganic molecules, RNA-precursors (brown symbols) as information carrier and the primordial RNA replicases or ribozymes, allowing them to eventually evolve into more efficient biological catalysts favouring corresponding PMBCs. The fluorescence image on the right shows an artificially assembled PMBC with encapsulated PMBCs based on prebiotic amphiphilic protein, visualized with a mEGFP domain. The enclosure of different sized molecules into our PMBC protocell model is shown in detail in Figure 3 (right column), illustrating the PMBC functionality as a potential dynamic reaction chamber. The dynamic nature of their membranes and fusion potential for the exchange of PMBC content is illustrated in Figure 4 (right column). Finally, the evolvement towards vesicles that undergoes the path towards cellular system with phospholipid bilayers as shown in today’s living cellular systems is illustrated. The principal compatibility of these two distinctive dynamic membrane systems is demonstrated in Figure 5 (right column) and presents the plausible transition of a prebiotic PMBC protocell model to current phospholipid membrane based cells. **b**, Illustrated are the different PMBC membrane compositions and encapsulated molecules investigated in this paper. Pictograms of the membrane constituents and enclosed molecules or amphiphilic proteins are denominated in Table 1. Different composition schemes are depicted in six segments surrounding the PMBC: There are three states with protein only membrane assembly His-H40I30, mEGFP-H40I30 and ComboRhod-K40I30 (10 to 16 o’clock) and three membrane states containing phospholipids or fatty acids with His-mEGFP-H40I30 or His-H40I30 (16 to 10 o’clock). Two PMBCs are depicted on the right, illustrating their functionalization through the encapsulation of catalytic reactions as step towards self-replicating systems. First we demonstrate the ability of anabolic biosynthesis processes by performing ligation reactions of small DNA oligonucleotides (15 bp) to large DNA (1200 bp) fragments within the PMBC described in Figures 6. Finally the implementation of in IVTT into the PMBC demonstrate the intrinsic ability of the PMBC protocells to synthesize their own structural constituents as basis for a self-replicable entity.

## Prebiotic peptides for PMBC assembly

There is agreement about a set of very abundant (Gly, Ala,Asp, Val) and less abundant prebiotic (aa) such as (Glu, Ser, Ile, Pro; His, Lys). The most abundant prebiotic aa G, A, D, V are spontaneously formed under Stanley Miller or similar prebiotic atmosphere conditions (endogenous sources) and were frequently found on meteorites or deep sea black smoker (exogenous sources)^29,21^. The latter prebiotic aa are structurally more complex and mainly evidenced from endogenous sources^21,30^. In our work, peptides composed of rather hydrophobic prebiotic aa such as Gly, Ile, Val, Pro and hydrophilic aa such as Ser, His and Lys have been applied (Table 1). Oligopeptides have been shown to be synthetically accessible under prebiotic conditions on clays^22^, minerals^23^, on metal and under hydrothermal conditions^31^ and even at air - water interfaces^32^. Schwendinger and Rode et al. showed the simplest mechanism of salt-induced peptide synthesis (SIPF) from prebiotic aa^33^. The consequent peptide chain elongation on clay minerals within the same environment and reaction conditions yields oligopeptides in reasonable amount and is applicable to all aa investigated so far (Gly, Ala, His, Val, Lys, Leu, Glu, Asp and Pro)^34^. Recently high yields up to 50 % and up to 20 aa could be synthetized applying ‘one-pot’ reaction conditions^35^. They demonstrated the co-condensation of glycine with eight other aa (Ala, Asp, Glu, His, Lys, Pro, Thr and Val), incorporating a range of side-chain functionality. Thus all the constituent amino acids and potentially the oligopeptides of the amphiphilic peptides employed in this work (Table 1) are amenable under prebiotic conditions.

**Table 1:**
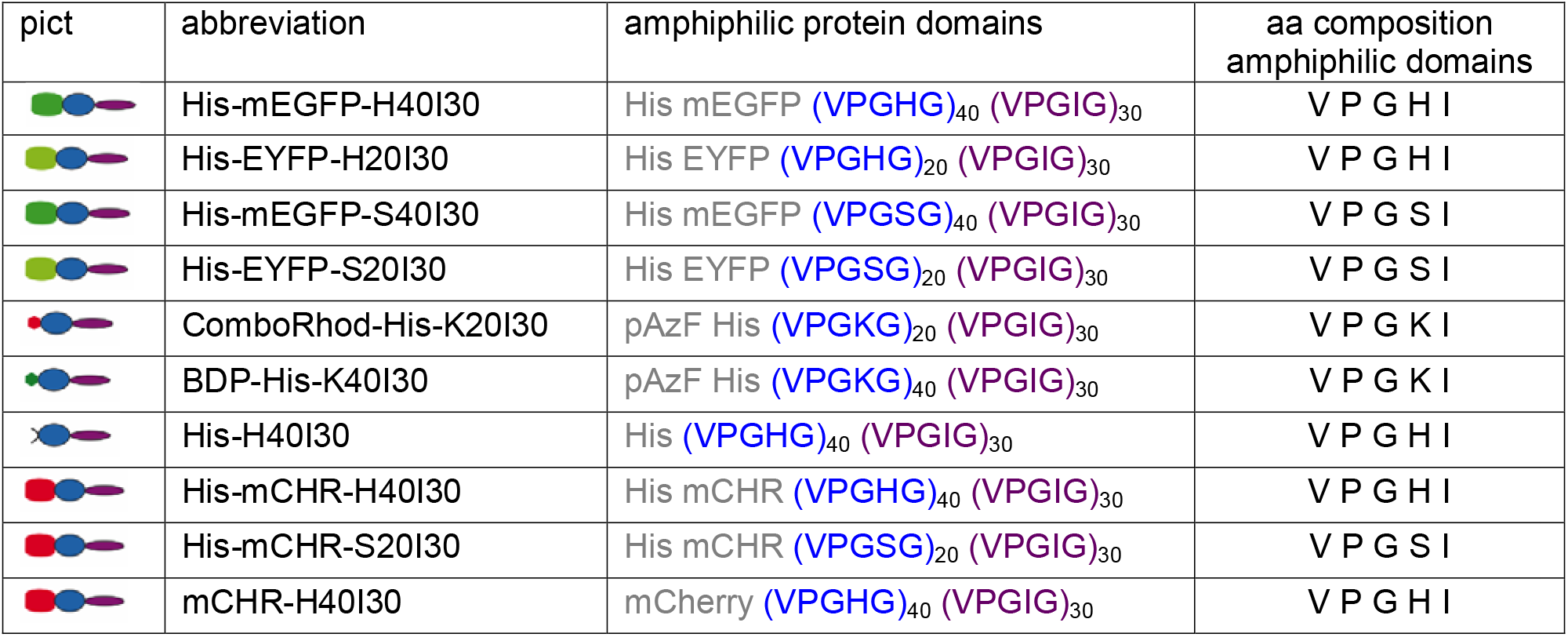
Amphiphilic protein block-copolymers for PMBC assembly. The first and second row specify the pictograms and abbreviations of the amphiphilic proteins used in this study. In the third row the amphiphilic protein blocks are listed. Grey letters mark the domains primarily responsible for purification (His) or visualization (mEGFP, EYFP, mCherry, pAzF) aspects. The blue (hydrophilic) and violet (hydrophobic) letters mark the protein domains responsible for the self-assembly property of the amphiphilic proteins. The repeat numbers of the pentameric peptide repetitions are indicated. The fourth row specify the amino acid components present in the amphiphilic protein domains. The correctness of the amino acid sequences of all applied ‘prebiotic’ amphiphilic proteins was confirmed by SDS PAGE and/or LC-MS/MS. (see Supplementary Data 1).

| pict | abbreviation | amphiphilic protein domains | aa composition<br>amphiphilic domains |
| --- | --- | --- | --- |
|  | His-mEGFP-H40I30 | His mEGFP (VPGHG)<sub>40</sub> (VPGIG)<sub>30</sub> | V P G H I |
|  | His-EYFP-H20I30 | His EYFP (VPGHG)<sub>20</sub> (VPGIG)<sub>30</sub> | V P G H I |
|  | His-mEGFP-S40I30 | His mEGFP (VPGSG)<sub>40</sub> (VPGIG)<sub>30</sub> | V P G S I |
|  | His-EYFP-S20I30 | His EYFP (VPGSG)<sub>20</sub> (VPGIG)<sub>30</sub> | V P G S I |
|  | ComboRhod-His-K20I30 | pAzF His (VPGKG)<sub>20</sub> (VPGIG)<sub>30</sub> | V P G K I |
|  | BDP-His-K40I30 | pAzF His (VPGKG)<sub>40</sub> (VPGIG)<sub>30</sub> | V P G K I |
|  | His-H40I30 | His (VPGHG)<sub>40</sub> (VPGIG)<sub>30</sub> | V P G H I |
|  | His-mCHR-H40I30 | His mCHR (VPGHG)<sub>40</sub> (VPGIG)<sub>30</sub> | V P G H I |
|  | His-mCHR-S20I30 | His mCHR (VPGSG)<sub>20</sub> (VPGIG)<sub>30</sub> | V P G S I |
|  | mCHR-H40I30 | mCherry (VPGHG)<sub>40</sub> (VPGIG)<sub>30</sub> | V P G H I |

The minimal amino acid and hence genetic sequence which allow to constitute amphiphilic proteins forming dynamic membranes require an efficient sequence motive which provides the structural requirements for dynamic membranes formation. The pentapeptide repeat motif applied in this work is derived from elastin like proteins (ELP) and based on the VPGVG repeat unit but is applied exemplarily for other simple amphipathic peptides with structure forming capability, amenable under prebiotic conditions. This ß-spiral repetitive sequence motif was applied due to its possible cylindrical geometry and dynamic analogy to phospholipids^36^ and their membrane forming capacities^37^.

In view of prebiotic peptide synthesis, repetitive peptide motifs comprise synthetic plausibility as first membrane constituents. In the presence of one abundant or dominant protein building block motif such as an individual prebiotic amino acid (e.g. glycine)^35^ or one prebiotic peptide motif (e.g. VPGHG) the multimerization of the motive (aa or peptide motive) has a high propensity. In addition, idealized computational model simulations predict a favored appearance of highly ordered and repetitive sequence patterns starting from monomer pools for autocatalytically active binary polymers^38^, supporting the plausible emergence of simple repetitive protein motifs. The role of peptides under early earth conditions is currently under debate in the context of the amyloid world hypothesis settling before the “RNA-world era”, postulating peptide amyloids as evolvable molecular entities able to self-replicate and transmit information^39^. In this context it was currently demonstrated that short repetitive peptide sequences may act as information coding molecules that catalytically allow the template-directed synthesis of defined repetitive peptide motives^39^. Whereas this was shown for repetitive amyloid ß-sheet motives, this could be a mechanism to receive and maintain sequence motives necessary for the formation of PMBCs. This demonstrates the bridging ability of replicating first information-coding molecules on the basis of prebiotic aa with the compartment forming capability.

The here presented pentapeptide repeat motif is known to tolerate mutations at the fourth position (valine; V) while maintaining its structural and functional properties^40^. This variability permits the adjustment of the protein domain properties by changing the number of repeat units or the amino acid at the fourth position. The hydrophilic protein domain was modified with hydrophilic prebiotic aa (S, K, H) at the fourth position of each pentapeptide motif and with hydrophobic prebiotic aa (isoleucine, I) at the same position to yield ‘prebiotic’ amphiphilic peptides (Table 1). N-terminal fusions of fluorescent proteins mEGFP and mCherry were added for direct characterization via fluorescence microscopy. The ratio of the amphiphilic block-domain proteins (Table 1) consisting of twenty or forty repeats of (VPGXG)_n_ (X= S,H,K) and thirty units of VPGIG were chosen due to geometric properties, hydropathic indices of the domains and stability results^41^. In addition, other hydrophilic (n=20) to hydrophobic ratios proved to be very efficient for vesicular assembly *in vivo* and *in vitro* by our group^37^. Even though the constituting amino acids and potentially the oligopeptides of the amphiphilic peptides employed in this work (Table 1) are amenable under prebiotic conditions, considerably shorter amphipathic peptides preserving the structure forming capacity would improve the feasibility of a prebiotic assembly scenario. The reduction of aa composition, complexity and length down to 9 pentapeptides proofed to preserve the vesicle forming capacity but resulted in poor assembly frequency and stability (see Supplementary Table 1 and Supplementary Fig. 1). The observed assembly of these short amphipathic peptides into PMBC emphasizes the plausibility of our hypothesis. However, complying the requirements regarding stability and assembly frequency, further PMBC characterization and applications in this work were conducted using the longer amphiphilic peptides (70 pentapeptides, Table 1). Successful PMBC assembly from different ‘prebiotic’ amphiphilic peptides is accomplished applying a very simple assembly method^42^ (see method section) (Fig. 2). The fluorescent images depict vesicles of 1 μm up to 15 μm enclosed by a homogenous fluorescent membrane boundary. Corresponding negatively stained TEM images show vesicular structures up to 800 nm in diameter. Amphiphilic peptides equipped with mEGFP or EYFP, different hydrophilic block length (n=20 vs n=40) and serine versus histidine as additional prebiotic aa were assembled (Fig. 2a and b). Successful PMBC assembly is demonstrated with two different hydrophilic block length (n=20 vs n=40) and prebiotic guest aa (H, S) (Fig. 2a, b) emphasizing the promiscuity of the composition of the amphiphilic peptides. In order to prove that PMBC assembly does not rely on ‘non prebiotic’ fluorescent proteins we introduced two different small fluorescent dyes (via click chemistry shown in Fig. 2c (for reaction details see Supplementary Fig. 2). Fluorescent images in red illustrate Combo-rhod-His-K20I30 and in green BDP-His-K40I30 assembled into PMBC of 0.7 up to 6 μm in diameter. This illustrates that successful PMBC formation does no rely on fluorescent fusion proteins. To also exclude the necessity of organic dyes for vesicular assembly, exclusively prebiotic His-H40I30 PMBCs were assembled and imaged via encapsulated dye (Fig. 2c right). This demonstrates the ability of PMBC assembly solely based on prebiotic aa. Two prebiotic PMBC assembly scenario are described below which yield vesicles with different efficiencies and composition. One method completely omitting any additional organic component is based on rehydration, while the second method applies alkanols yielding PMBC in very high abundance.

**Figure 2:**
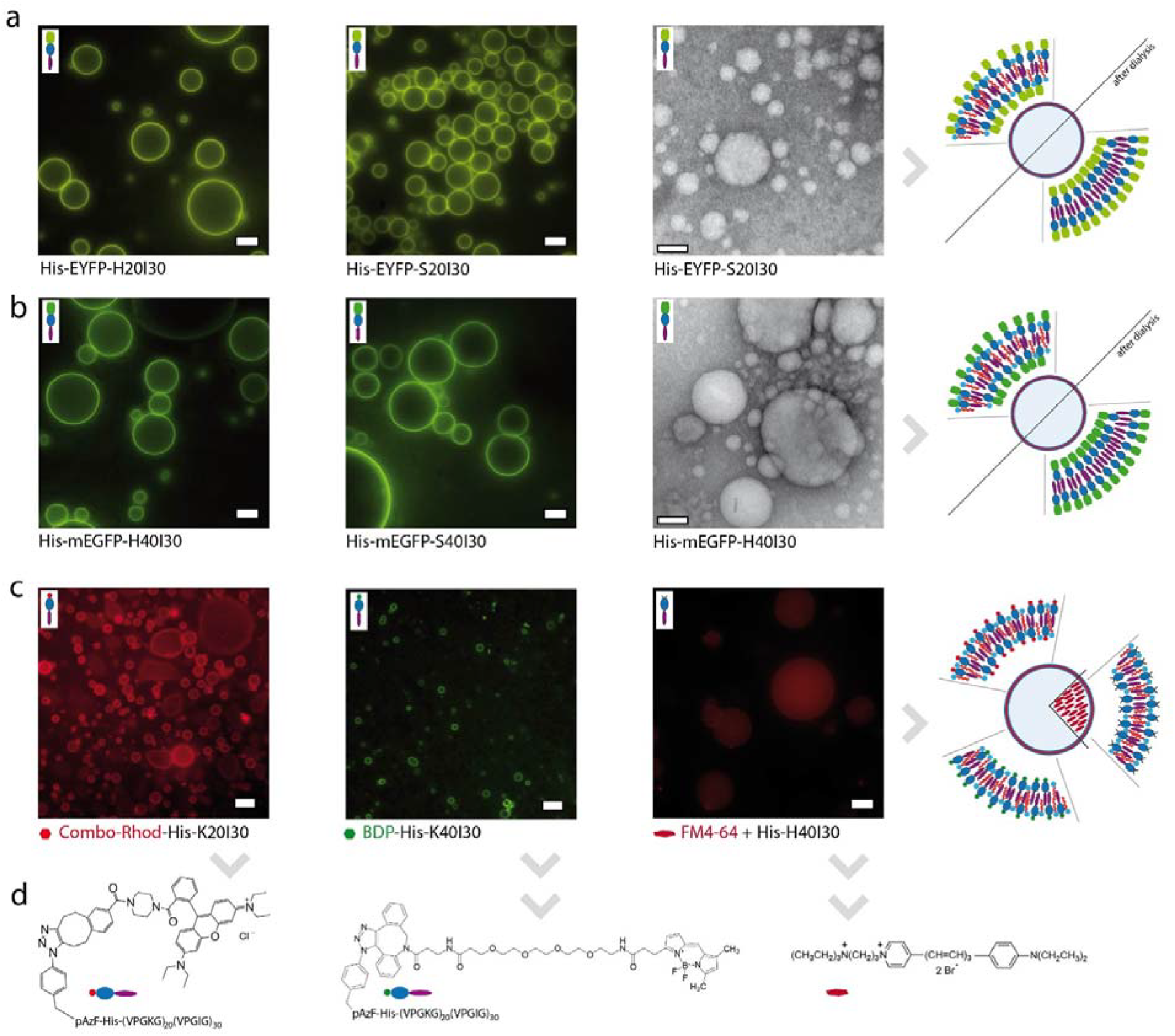
Epifluorescence images and TEM images demonstrate the formation of protein membrane based compartments (PMBC) from amphiphilic ELP building blocks composed of prebiotic aa. **a**, Epifluorescence images of PMBCs assembled of amphiphilic proteins composed of prebiotic aa which are supplemented with an EYFP domain for visualization. All mentioned constructs in this Figure have a (VPGIG)_30_ domain as the hydrophobic block of the amphiphilic protein. The left image shows an amphiphilic ELP with the hydrophilic histidine ELP domain (VPGHG)_20_ and an additional EYFP domain. The middle image shows a correspondent amphiphilic protein with a hydrophilic serine ELP domain (VPGSG)_20_ instead of histidine. The right image presents a negatively stained TEM image of the same construct as shown in the middle image. The illustration at the right end present the membrane composition of the PMBC constituted from EYFP supplemented amphiphilic proteins and n-butanol before and after dialysis. **b**, Images of PMBC assembled of amphiphilic proteins composed of prebiotic aa supplemented with a mEGFP domain for visualization in epifluorescence microscopy. The left image shows an amphiphilic ELP with the hydrophilic histidine ELP domain (VPGHG)_40_ and the mEGFP domain. The middle image shows a correspondent amphiphilic protein with a hydrophilic serine ELP domain (VPGSG)_20_ instead of histidine. The right image present a negatively stained TEM image of the same construct as shown in the left image. The illustration at the right end present the membrane composition of the PMBC constituted from mEGFP, supplemented amphiphilic proteins and n-butanol before and after dialysis. **c**, Images of PMBCs built of amphiphilic ELPs, supplemented with the unnatural amino acid p-azido-phenylalanine (pAzF) which is labelled with a chemical dye as indicated combo-rhodamine (left), DBCO - bodipy (middle) (see methods and SI Fig 1). In the right image the chemical dye (FM4-64) is encapsulated within the assembled compartments composed of a prebiotic amphiphilic ELPs without a fluorescent domain. The hydrophilic domains (from left to right) are (VPGKG)_20_, (VPGKG)_40_ and (VPGHG)_40_. The illustration at the right end present the membrane composition of the PMBC constituted from the amphiphilic proteins in the left images, n-butanol and the encapsulation of the FM4-64 dye. **d**, The formulas of the chemical dyes used in d) are presented and illustrated with respective pictograms. The scale bars correspond to 5 μm in epifluorescence images and 100 nm in TEM images.

Simple dehydration of the amphiphilic protein film on glass slides and rehydration with subsequent swelling of amphiphilic protein film yields vesicles up to 2 μm (Supplementary Video 1). This very simple vesicular self-assembly only requires prebiotic amphiphilic proteins, water and one cycle of de- and rehydration to yield PMBCs, illustrating a plausible prebiotic assembly scenario. In the second method, ‘prebiotic’ amphiphilic proteins are extruded with 5% to 10% v/v alkanols such as n-butanol or 1-octanol to induce the PMBC self-assembly with high yield of PMBCs up to 15 μm in diameter. Illustrating the robustness and prebiotic relevance, the salt concentration during assembly may vary from zero up to 5 M and the temperature during vesicle formation can range from 4°C up to 70°C. This assembly method is applied throughout all experiments if not stated otherwise and details can be found in the methods section. After dialysis against buffer and removal of n-butanol the PMBC are stable up to 100°C, pH 2- 11 and at least 8 month storage at room temperature (Supplementary Fig. 3). When compared to the alkanol-free swelling protocol, the number of vesicles is roughly increased a thousand fold. The very simple protocol of mixing amphiphilic proteins with fatty acid derivatives (fatty alcohols) is also of prebiotic relevance, since fatty acids and fatty alcohols have been synthesized at prebiotic conditions^26,27^ and found on meteorites in high abundancies^28^. Fatty acids and their corresponding alcohols are attractive candidates for protocell constituents itself^43^. They form bilayer membrane vesicles^28^ that are capable of growth and division^44,45^ and such membranes are highly dynamic^6^. So far all proposed prebiotic protocell boundaries are based on pure and single constituents^2^, even though prebiotic reactions conditions and meteorites contain always a mixture of molecules of different classes (aa, fatty acids, sugars). In addition, recent studies have shown, that mixtures of amphiphiles allow for an efficient and more stable formation of membranes^46^. Therefore we further tested compatibility of our PMBC by extruding simple fatty acids (e.g. lauric acid) together with our prebiotic amphiphilic proteins also yielding PMBC in medium concentrations (Supplementary Fig. 3). The compatibility of alkanols and fatty acids to our PMBC increases the plausibility of a mixed assembly scenario even though PMBC assembly solely necessitates prebiotic amphiphilic proteins as described above (Supplementary Video 1). However when compared to pure fatty acids protocell models^6^ the assembly protocol and PMBC stability is far more robust under different environmental conditions when compared to the very promising fatty acids protocell model^43^ which are stable only in a narrow pH range and low ionic strength^47^. In addition, vesicular assembly of PMBCs requires an amphiphilic protein concentration four orders in magnitude less compared to lipid vesicles (10 mM fatty acids^6^ vs 1 μM of amphiphilic protein) and tolerates up to 5 M salt which is contrast to fatty acid vesicles. Hence, given the successful synthesis of amphiphilic protein building blocks, the formation of protocell-like compartments at such low concentration may have interesting advantages in a prebiotic world. In the light of environmental conditions on the prebiotic earth, the stability of PMBC up to 100°C and pH ranges from 2- 11, supports the plausibility of prebiotic PMBC as first protocell models.

## Encapsulation of molecules into PMBCs

The encapsulation of intact molecules into PMBCs is one important step towards the design of artificial cells and characterizes the PMBC properties and membrane permeability. The addition of molecules to be encapsulated during PMBC assembly leads to the inclusion of these molecules into the vesicular lumen (Fig. 3). During assembly these PMBCs can encapsulate molecules of different length scales, ranging from small dyes (800 Da) to macromolecular dyes (3 kDa), to functional completely folded proteins (28 kDa) and PMBCs (8 μM) themselves (Fig. 3a-c). This illustrates a complete or partial membrane tightness and diffusion barrier towards molecules of different nature such as sugars (dextran), positively charged small molecules (ATTO Rho 13) and completely folded proteins, essential under harsh environmental conditions. Further these results demonstrate that this assembly protocol does not compromise the functionality of the encapsulated dyes or proteins. In addition, the internalization of whole PMBCs inside another PMBC (Fig. 3b) illustrates the possibility of sub compartmentalization of spatially separated reactions within one prebiotic reaction compartment (Supplementary Video 2). Another requirement for prebiotic protocells, apart from molecular encapsulation, is membrane fluidity, fusion and fission potential which is described below (Fig.4).

**Figure 3:**
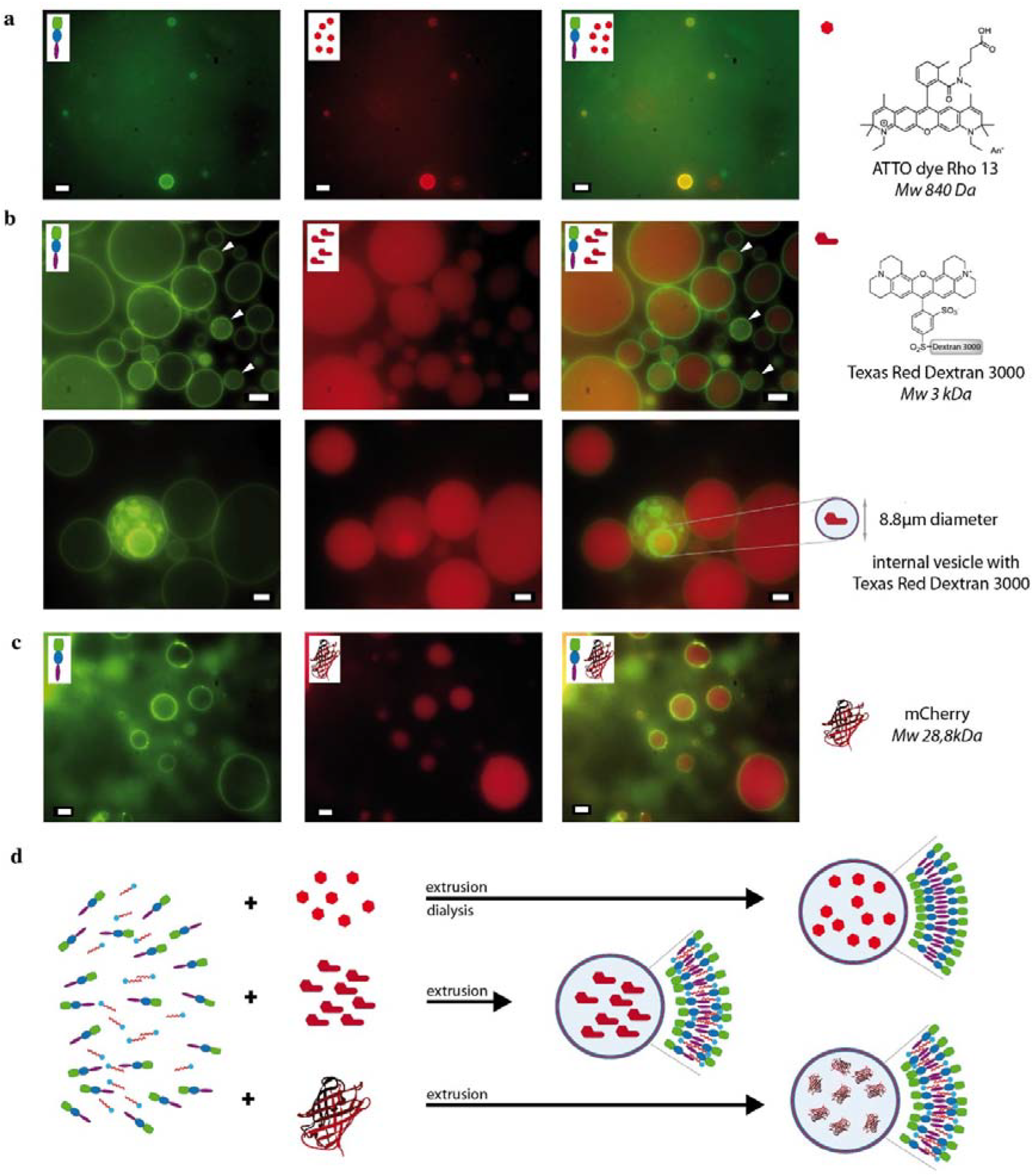
PMBC encapsulate small dyes, macromolecular dyes, functional proteins and small PMBCs as premise for functional reaction chambers. **a**, Fluorescence images in green (left), red (middle) and merged channel (right) demonstrate that a PMBC assembled from His-mEGFP-H40I30 and n-butanol encapsulate small ATTO Rho 13 dye during assembly. Subsequent dialysis after assembly against 10 mM Tris-HCl buffer pH 8 leads to removal of n-butanol and excess of ATTO Rho 13. **b**, Fluorescence images (upper and lower row) in green (left), red (middle) and merged channel (right) demonstrate that a PMBC assembled from His-mEGFP-H40I30 and n-butanol encapsulates macromolecular Texas Red Dextran3000 dye during assembly. The encapsulated dye concentration between PMBC differs (marked with white arrows in left and right upper image). The lower images demonstrate the encapsulation of smaller PMBC (containing the red dye) and protein membrane fragments within the PMBC assembly process. **c**, Fluorescence images in green (left) red (middle) and merged channel (right) demonstrate that a PMBC assembled from His-mEGFP-H40I30 and n-butanol encapsulate correctly folded mCherry during assembly. On the right of each row (a, b, c) formula and pictogram of the enclosure is illustrated together with their mass or dimension. Pictograms in the left upper edge of each image illustrate the visual constituents of the reaction. All scale bars correspond to 5 μm. **d**, Pictograms illustrate the process of assembly and encapsulation of chemical dyes and mCherry into the PMBC. In case of extrusion without subsequent dialysis residual amounts of n-butanol could be present within the dynamic protein membranes as illustrated.

## PMBC membrane dynamics

Dynamic membrane properties are required for membrane growth, exchange of vesicular contents and nutrient intake, e.g. exo- and endocytsosis^5^. Membrane fusion is a process important to many cellular events, especially for transport of nutrients or signaling. The dynamic properties and fluidity of the protein based membrane is shown in Figure 4. The fusion of cell sized PMBCs composed of BDP-His-K40I30 is illustrated in Figure 4a and b and Supplementary Video 3. PMBC fusion events are observed on distinct areas of the microscope cover slide, where PMBCs are adherent on the glass slide. In Figure 4 two fusing adherent 10 μm sized PMBC can be observed whereas in Figure 4b only one of the two fusing PMBCs is adherent. Surface adhesion of vesicles often leads to tensile membrane stress, flattening and deformation^48^ which can be observed in Figure 4a and b. The resulting increase in interfacial adhesion energy leads to membrane stretching and the exposure of hydrophobic areas and pockets and provokes the ultimate breakthrough fusion event^36^. In addition, PMBCs composed of His-mEGFP-H40I30 in buffer lacking alkanols undergo fusion events observed on the TEM grid surface (Fig. 4c), illustrating that alkanols are not required for fusion. In order to demonstrate fusion events and phase separation of assembled PMBC fusion in solution, two variants of ‘prebiotic’ amphiphilic proteins were equipped with two different fluorescent fusion proteins each. The right images in Figure 4e and f illustrate PMBC membrane patches of different color (red and yellow or green) belonging to a single PMBC. These patches indicate a previous fusion event of PMBCs equipped with different fluorophores (mCherry versus EYFP or mEGFP) but composed of the same amphiphilic domains H40I30 and S20I30 respectively. This indicates that round shaped PMBC fusion events in solution at RT does occur and that fusing membranes stay phase separated for at least 20 min. However, mixing of PMBC constituted of different ‘prebiotic’ amphiphilic proteins (His-mEGFP-S40I30 and His-mCherry-S20I30), here with a substantial length difference in the hydrophilic domain does not lead to visible fusion events (Supplementary Fig. 5). This observation indicates a specific or selective type of fusion where protein membranes composed of the very same constituents selectively fuse with each other. Such fusion events are frequent in contemporary phospholipid bounded cells where membrane proteins e.g. at synapses usually initiate specific fusion events^49^. In the protocellular context the dynamic PMBC membrane permits specific fusion and therefore the potential for directed exchange of vesicular content for e.g. more complex multistep reactions or exchange of evolved ribozymes. Another important property for prebiotic PMBCs – as new type of protocell model is the compatibility with contemporary living systems.

**Figure 4:**
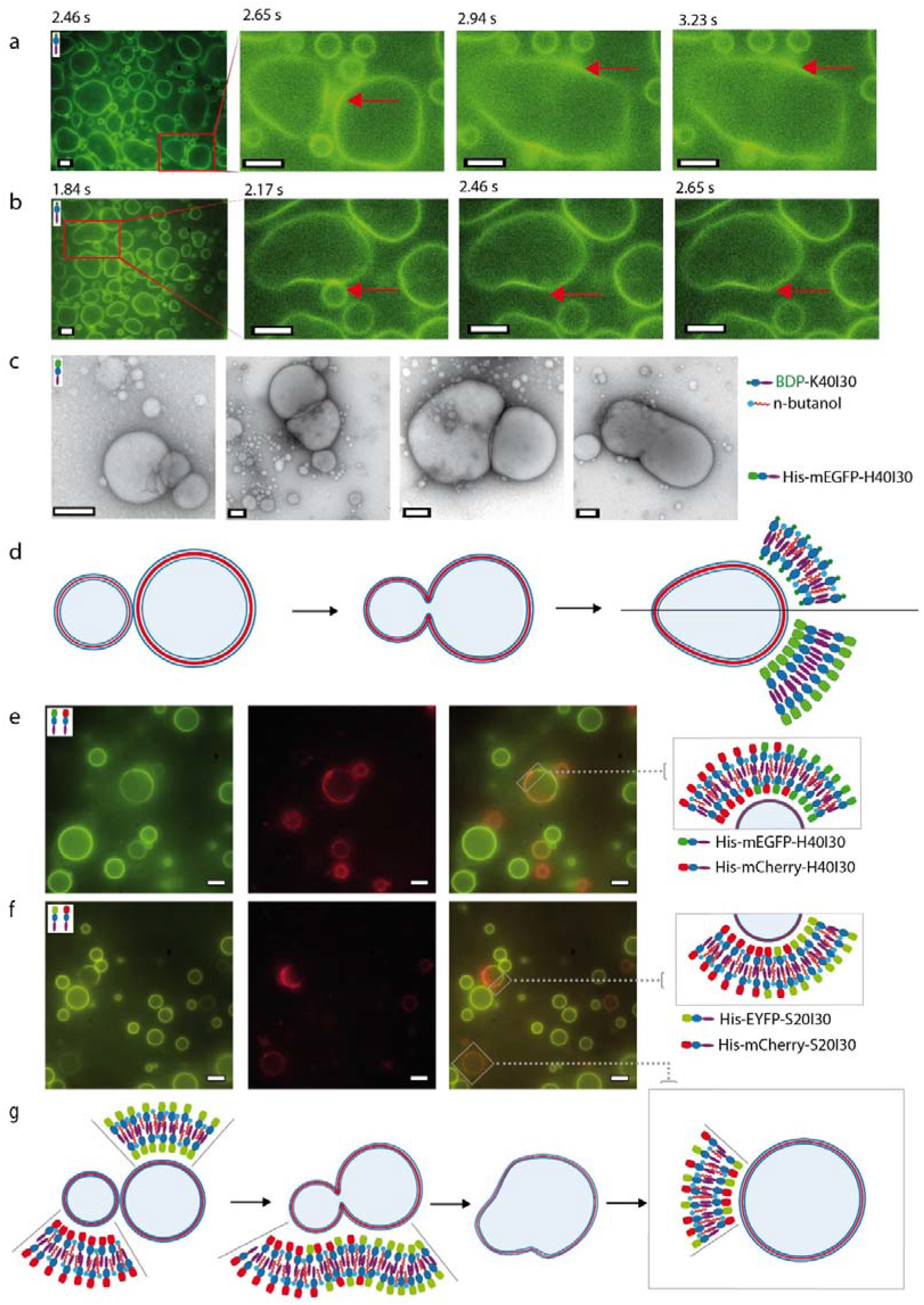
Epifluorescence images demonstrate the fusion of cell sized PMBC assembled from different amphiphilic proteins. **a**, Time series of the fusion of two 10 μm sized PMBCs consisting of BDP-K40I30 in 10 mM Tris-HCl pH 8 and 10% v/v n-butanol recorded with 4 frames per second in the green channel. **b**, Time series of the fusion of a 3 μm and 14 μm sized PMBC consisting of BDP-K40I30 in 10 mM Tris-HCL pH8 and 10% v/v n-butanol recorded with 4 frames per second in the green channel. **c**, Negatively stained TEM images of the fusion of 150 nm (left) up to 700 nm sized PMBC (right) consisting of His-mEGFP-H40I30 in 10 mM Tris-HCL pH 8.**d**, Schematic illustration of the fusion of PMBC assembled from amphiphilic proteins (a to c). **e**, Epifluorescence images of post assembly mixtures of two variants (His-mEGFP-H40I30 and His-mCherry-H40I30) of PMBCs in 10mM Tris-HCl pH 8 and 10% v/v 1-octanol. The left image in the green channel visualizes His-mEGFP-H40I30 proteins and PMBCs assembled thereof. In the middle image the red channel visualizes the His-mCherry-H40I30 proteins and PMBCs assembled thereof. The right epifluorescence image displays an overlay of the green and red channel images and reveals disjoined PMBCs and also fused PMBCs consisting of amphiphilic proteins with both fluorophores (mEGFP and mCherry). The grey rectangle marks a fused PMBC and is illustrated with pictograms of the present amphiphilic proteins at the right of this image. **f.** Epifluorescence images of two PMBC variants (one composed of His-EYFP-S20I30 and the other His-mCherry-S20I30) in 10mM Tris-HCl pH 8 and 10 % v/v 1-octanol. In the left image the green channel visualize the His-EYFP-S20I30 proteins and PMBC assembled thereof. In the middle image the red channel visualizes the His-mCherry-S20I30 proteins and PMBCs assembled thereof. The right image displays an overlay of the green and red channel images and reveal a disjoined PMBC and also a fused PMBC consisting of amphiphilic proteins with both fluorophores (EYFP and mCherry). The grey rectangle marks a fused PMBC and is illustrated with pictograms of the present amphiphilic proteins at the right of this image. **g**, Schematic illustration of the fusion process of PMBCs consisting of variants of amphiphilic proteins (here His-EYFP-S20I30 and His-mCherry-S20I30). The quadratic rectangle in the right image gives an example where the amphiphilic constituents are distributed homogeneously within the resulting PMBC membrane after fusing. Pictograms in the upper left edge of each image illustrate the PMBC constituting amphiphilic proteins. Scale bars of the epifluorescence images correspond to a 5 μm and scale bars in TEM images correspond to 200 nm.

## Liposome and phospholipid compatibility to PMBCs

The main building block of extant cellular membranes are phospholipid bilayers. In Figure 5 we show the compatibility of PMBC with one of the main contemporary cell membrane constituents. Multilamellar liposomes (red) based on 1,2-dioleoyl-sn-glycero-3-phosphoethanolamine (DOPE) can fuse and subsequently be internalized into PMBCs just by simple mixing (Fig. 5a and b). At low concentrations DOPE ATTO red dye 647N (1,2-dioleoyl-sn-glycero-3-phosphoethanolamine) is homogeneously incorporated into protein based vesicle membranes (green) upon swelling of vacuum dried PMBCs (Fig. 5d and e). At low concentrations DOPE lipids associate into the protein membrane whereas DOPE assembled in liposomes favours the intermolecular interaction with their analogues instead of protein amphiphiles (Fig. 5a). This process can be described as sort of phase separation in analogy to lipid-lipid phases separation. Such phase separations are due to collective properties of macromolecules and particularly relevant for the spatial organization of metabolic reactions in living organisms^50^. In our previous work^37^, we observed a strong interaction of non ‘prebiotic’ PMBCs with the lipid based cellular membrane of *E.coli, in vivo.* The close association and invagination of the cellular membrane associated with PMBC was observed using TIRF structured illumination microscopy. Those PMBCs where assembled from peptides containing some non-prebiotic aa (e.g. Glu). However the association of the *E.coli* membrane and PMBCs also demonstrates a strong interaction and fusion tendency of lipid bilayers and dynamic protein membranes *in vivo*^37^. The compatibility of phospholipids and lipid membranes with PMBC demonstrates the plausible transition of a prebiotic PMBC protocell model to current phospholipid membrane based cells.

**Figure 5:**
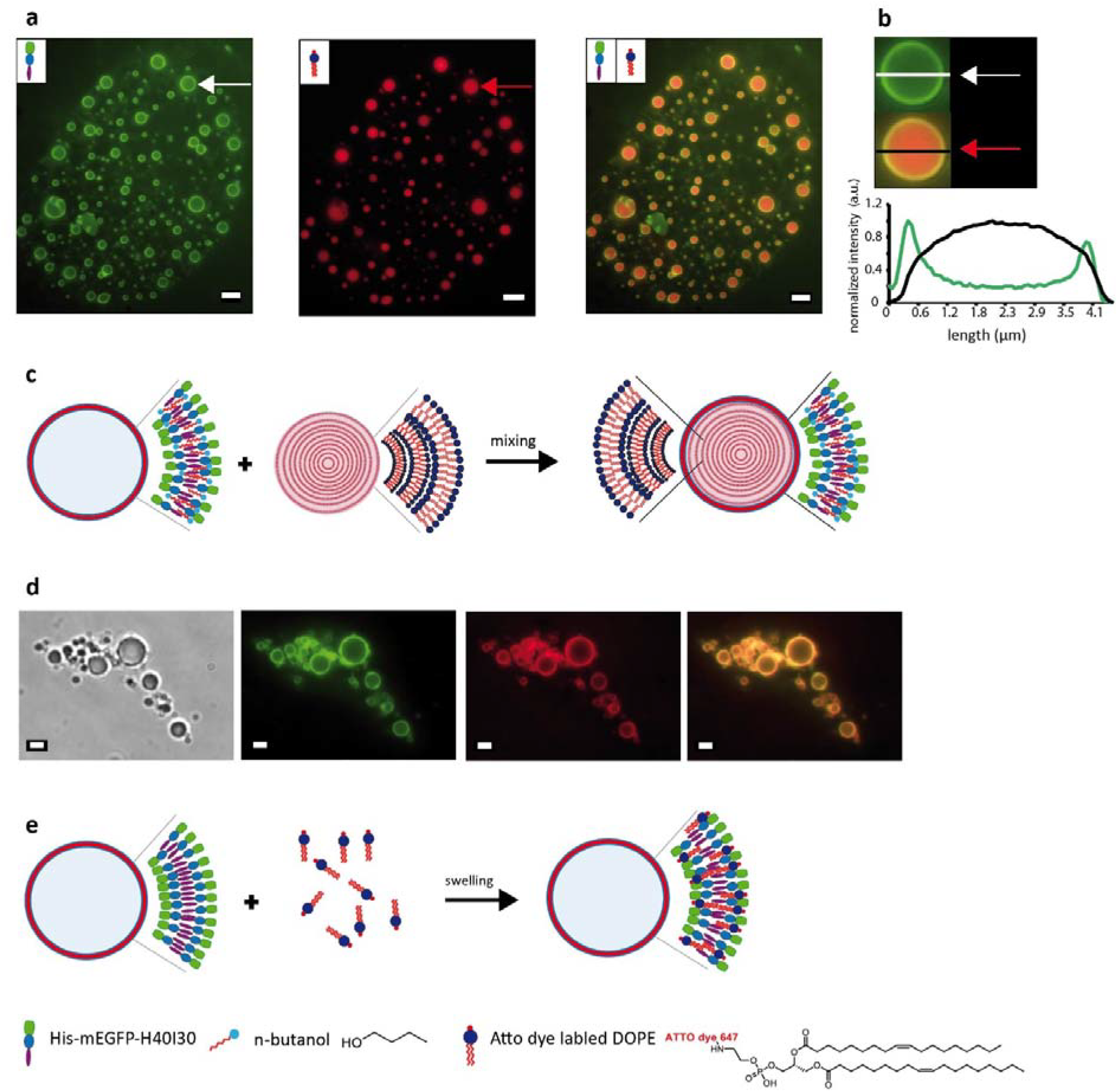
Epifluorescence images demonstrate the fusion with and subsequent internalization of contemporary phospholipids (red) by ‘prebiotic’ PMBCs (green). Epifluorescence image of **a**, His-mEGFP-H40I30 based PMBC in 10 mM Tris-HCL pH 8 and 10% v/v n-butanol in green channel (left), multilamellar DOPE (1,2-dioleoyl-sn-glycero-3-phosphoethanolamine) based liposomes in red channel (middle) and merged channels (right) demonstrate successful incorporation of DOPE-based liposomes into PMBC (green) **b**, Line scans of left and right images illustrate the successful fusion of multilamellar DOPE-based liposomes (red) with PMBCs. **c**, Schematic illustration of the fusion of DOPE-based liposomes and PMBC-based on His-mEGFP-H40I30. **d**, Whitefield (left) and epifluorescence images demonstrate the homogeneous incorporation of DOPE ATTO red dye 647N (1,2-dioleoyl-sn-glycero-3-phosphoethanolamine) (red channel) into protein based vesicles membrane (green) upon mixing with vacuum dried PMBCs. The merged green and red channel on the right demonstrate the homogenous incorporation. **e**, Schematic illustration of the internalization of DOPE ATTO red dye 647N into the protein membrane. The scale bars correspond to 5 μm in a) and 2 μm in d).

## Anabolic reaction pathways towards self-sustaining PMBC protocells

The simple encapsulation of functional molecules into the PMBC was shown in Figure 3 and is an important requirement in building artificial cells. However, a major step towards a self-sustaining protocell model is the capability to assemble and propagate genetic information and the ability to execute anabolic reactions as e.g. the polymerization of RNA, DNA molecules or aa. In Figure 6 we present the ligation of 15 base pairs (bp) oligonucleotides (encoding for one pentapeptide ELP motive) to high molecular DNA strands with up to 1200 bp within our prebiotic PMBCs. This demonstrates the ability of PMBCs to embed the polymerization of genetic information that subsequently could serve as an evolvable template polymer for the repetitive protein membrane constituents of the PMBC. The length of resulting ligated DNA potentially encodes for proteins of up to 80 pentapeptide repeats which fits well to the DNA template size of our investigated amphiphilic PMBC constituents. All ligation reactants were encapsulated during PMBC assembly. After 20 h DNA ligated outside the PMBC was captured via silica particles modified for DNA capture (Fig. 6a, I.)

**Figure 6:**
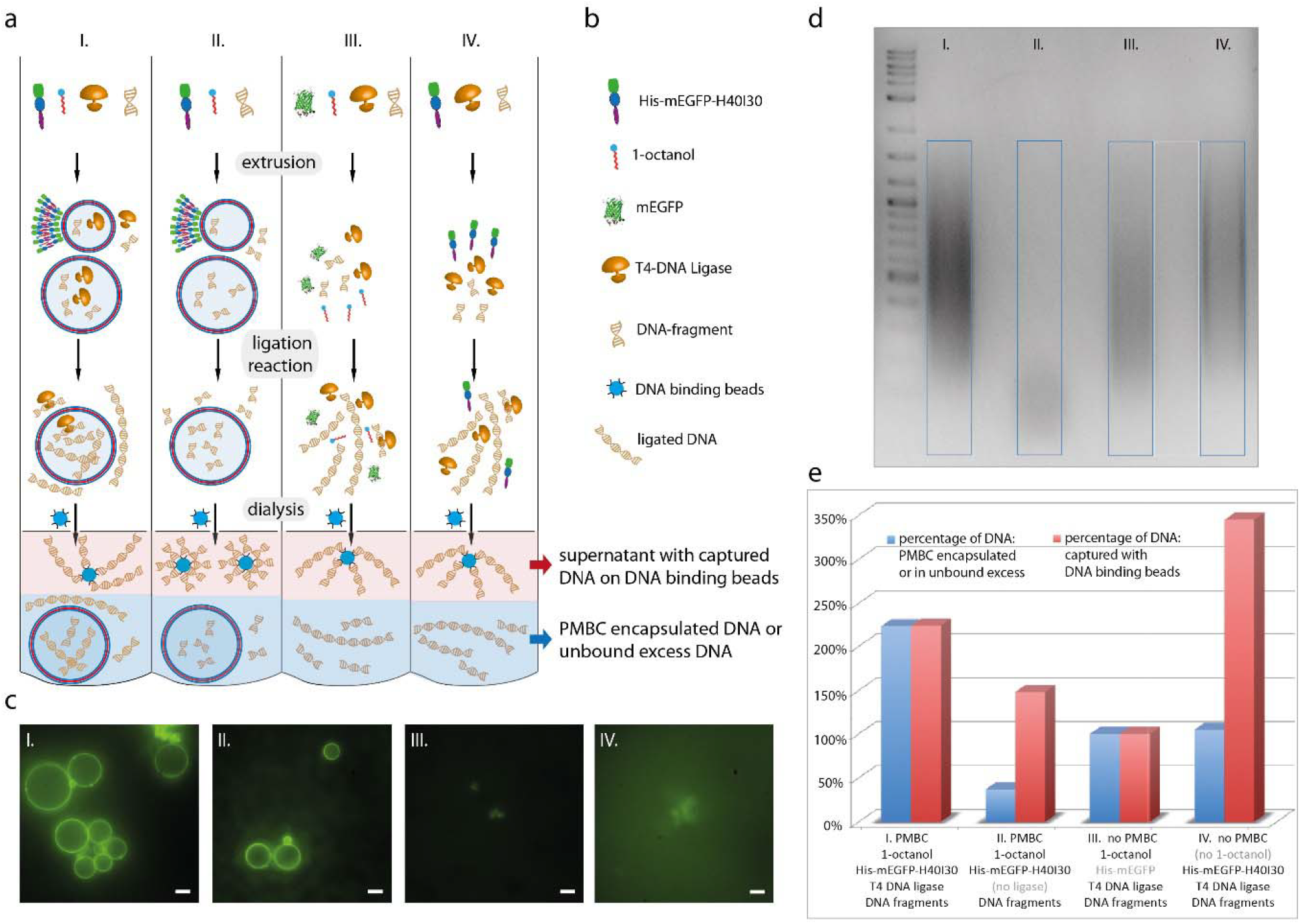
Successful ligation of short DNA oligos (15 bp) to high molecular weight DNA (1200 bp) inside prebiotic PMBC. **a**, Illustrative flowchart depicts the ingredients, the experimental approach of the four variants (I. to IV.) of ligase and control reactions. The light red and blue fields at the bottom of each reaction process mark the purified reaction products used for quantitative analysis via agarose gel-electrophoresis. The red field and the red arrow indicate the DNA captured with DNA binding beads (Supplementary Fig. 6). The blue field and the blue arrow indicate for the ligation products uncaptured because of PMBC encapsulation or in excess with respect to the DNA binding beads binding capacity. These fractions (I. to IV.) are separated on a 1.5% agarose gel (see c) for quantitative analysis. **b**, Annotation of pictograms used in a). **c**, Epifluorescence images of an aliquot of the dialyzed reaction volumes (I. - IV.) demonstrate PMBC formation in reactions I. and II. and no PMBC formation in III. and IV. The scale bars correspond to 5 μm. **d**, The DNA products of the four described (ligation-) reactions (sheltered or unbound fraction: blue field in a) are separated and visualized on an agarose gel. The probe volume in each lane is proportional to original DNA fragment amount of each reaction. Blue squares mark the regions of interest used for quantification of the amount and distribution of DNA ligation products (presented in e) by measuring the grey values compared to the background (white square) grey values with ImageJ analysis software. **e)** Quantitative diagram of the percentage amount of sheltered and unbound DNA in the four reactions (I. to IV.). Control III. is taken as 100% reaction reference. Here no PMBCs are formed but all reaction components (T4-DNA-Ligase, 1-octanol, protein (mEGFP) and short DNA-fragments) necessary for ligation reaction are present, whereas the PMBC forming protein His-mEGFP-H40I30 is replaced with mEGFP. The ligation experiment was conducted in at least three independent assays. The column heights are depicted relative to III. Blue bars indicate for the encapsulated or unbound DNA fractions (see c, and a, blue fields). Red bars correspond to the captured DNA (see Supplementary Fig. 6 and a), red field).

To prove successful ligation within the PMBC three different control ligations were performed. First, we omitted the T4 DNA Ligase (control II, Fig. 6a, II.) second, we exchanged the PMBC constituent against mEGFP domain that is not able to form PMBCs (control III, Fig. 6a, III.) and finally 1-octanol (control IV, Fig. 6a, IV.) was omitted to prevent PMBCs formation. The reaction process is illustrated in Figure 6a by a flowchart of schematic pictograms explained in Figure 6b. Subsequently to the ligation reaction of I. to IV, reaction volumes were dialyzed and the presence or absence of PMBC within these reactions are visualized with epifluorescence microscopy (Fig. 6c). The large fraction of DNA for control II-IV in the supernatant (Fig. 6a, I., red fields) was captured through adding equal aliquots of DNA binding silica particles to each reaction format and analyzed via an agarose gel (Supplementary Fig. 6) The remaining DNA, which was in excess and not bound to particles (for control II-IV) or sheltered within the PMBC was analyzed and quantified via agarose gel electrophoresis (Fig. 6d and e blue bars). Quantitative analysis of the grey values of the corresponding gel lanes (for details see methods section) demonstrates clearly that in comparison to the control reactions (control III corresponds to 100%) the amount of analyzed DNA is significantly increased from supernatant captured DNA towards sheltered uncaptured DNA (Fig. 6. I. versus control IV). Consequently, for ligation reaction I an equal amount of DNA ligation (Fig. 6a - e) was performed inside the assembled PMBCs versus outside. Therefore, we can conclude that anabolic reactions such as the polymerization of genetic information could be conducted within PMBC realizing an important step towards a self-sustaining protocell model.

Taking one definition of life as “a self - sustaining chemical system capable of Darwinian evolution^51^ the ability for self-replication is crucial for valid protocellular models. In Figure 7 we demonstrate the DNA encoded synthesis and subsequent incorporation of protein-based membrane constituents through IVTT within the PMBC as essential step towards a self-replicable system. For this experiment, preassembled and vacuum dried PMBCs were swollen on glass slides together with IVTT mix in buffer. The PMBC swelling leads to partial encapsulation of IVTT mix containing the DNA which encodes the membrane constituent mCherry-H40I30 and leads to its subsequent expression and membrane incorporation. The resulting red membrane fluorescence signal was imaged (Fig. 7b) after 5 hours of expression and quantified through line plots across the PMBC profile (Fig. 7c and d). In order to distinguish between out and inside expression, three IVTT conditions depending on suppression location (for details see method section) where defined: IVTT outside and inside (no suppression) (I), IVTT inside (suppression outside) (II), and no IVTT (suppression in- and outside) (III). Here kanamycin (Kan) as strong prokaryotic ribosomal translation inhibitor was applied for IVTT inhibition and was comparable to chloramphenicol, RNAase or DNAse induced inhibition (data not shown). For condition I (IVTT outside and inside) the strongest red membrane fluorescence and greatest SmN and SmL ratios (defined as in Fig. 7c right panel) can be observed in Figure 7c and d (I) (see Supplementary Table 3). For the desired scenario (IVTT inside and outer suppression) a significantly smaller SmN and SmL ratio is measured. This significant lower red membrane fluorescence (compared to condition I) indicates that mCherry-H40I30 is exclusively expressed inside the PMBC and gets incorporated into the PMBC membrane from inside. A difference in red membrane fluorescence signal intensity of single PMBC for condition II is observed in Figure 7b II (multiple PMBC within the red image). This variation can be associated with variable IVTT mix encapsulation efficiency. High variations of encapsulation efficiencies for the applied swelling method are described in the literature for lipid based vesicles^52,10^. The suppression of IVTT outside and inside the PMBC leads to weak red auto fluorescent background signal where the red signal intensity of lumen and vesicle membrane are indistinguishable (SmL ratio = 1.1) (line plot of Fig. 7c III). This can be attributed to Kan induced complete suppression of mCherry-H40I30 expression and the lacking membrane incorporation. The measurement of the red fluorescence of the entire sample solution confirms a significantly higher signal of IVTT inside compared to the background signal for the no IVTT condition (Fig. 7d (i) and Supplementary Table 4). Vogele et al.^53^ recently obtained very similar results for IVTT inside ELP-based vesicles constituted from non-prebiotic (VPGEG)n and (VPGFG) ELP domain derivatives previously described to assemble and to be modified chemically *in vivo* and *in vitro* by our group^37,42^. This demonstrates the feasibility of IVTT inside dynamic PMBC independent of the exact molecular composition of the membrane building blocks. Applying the IVTT of prebiotic membrane building blocks we could demonstrate an important step towards a self-replicating prebiotic PMBC protocell via DNA encoded synthesis of protein-based amphiphilic membrane constituents and their subsequent insertion into the prebiotic PMBC membrane.

**Figure 7:**
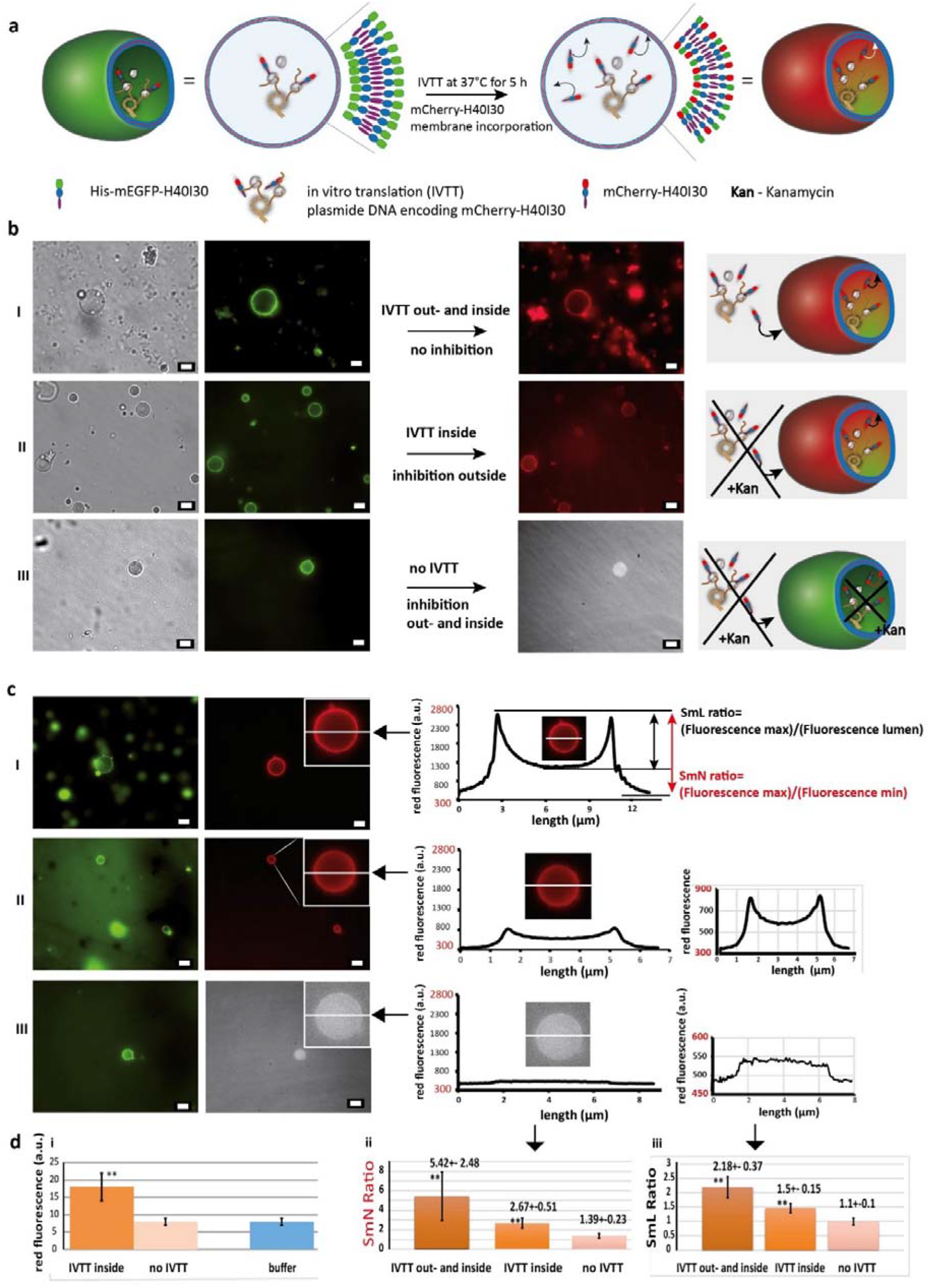
A step toward self-replicating prebiotic PMBC protocells: Synthesis and incorporation of the PMBC membrane building block mCherry-H40I30 (red) inside PMBC (assembled of His-mEGFP-H40I30) via *in vitro* transcription and translation (IVTT). **a**, Schematic illustration of a PMBC assembled from His-mEGFP-H40I30 (green) after encapsulation of the IVTT mix, (see methods section) and plasmid DNA encoding for mCherry-H40I30. After encapsulation of the IVTT mix into the PMBC upon swelling on glass slides, mCherry-H40I30 expression and subsequent membrane incorporation leads to red fluorescent PMBC. **b**, White field, green and red fluorescence images of PMBCs after 5 hours of IVTT with three different IVTT conditions. **(I)** In the absence of kanamycin (Kan) the IVTT of mCherry-H40I30 inside and outside of the PMBC leads to strong red fluorescent membrane signal and red protein agglomerates outside. **(II)**The addition of Kan to the outer PMBC solution inhibits outer IVTT (no red agglomerates) and leads to a red fluorescent membrane signal of the PMBC via IVTT and incorporation of mCherry-H40I30 from inside the PMBC. The different red membrane intensities demonstrate the variable IVTT mix encapsulation efficiencies upon swelling. **(III)** The addition of Kan to the outer and inner PMBC solution entirely inhibits IVTT outside and inside the PMBC. The weak red signal results from red auto fluorescence of the PMBC upon excitation at 587 nm. **c**, Green and red fluorescence images of PMBCs after 5 hours of IVTT of mCherry-H40I30. **(I)** Line plots of the red fluorescent PMBC upon IVTT inside and outside the PMBC show a 5.4 ± 2.5 fold membrane signal to noise ratio (SmN ratio) and 2.2+/-2.2 fold membrane signal to lumen ratio (SmL) (also see d (ii, iii)).**(II)** Line plots of the red fluorescent PMBC upon IVTT inside the PMBC show a 2.7+-0.5 fold SmN ratio and about 1.5 +-0.15 fold SmL ratio (also see d (ii, iii)). This indicates that about half (SmN ratio 2.7 versus 5.4) of the expressed mCherry-H40I30 was synthesized and incorporated into the PMBC membrane inside the PMBC when compared to condition (I). **(III)** Line plots of the control PMBC (no IVTT) where mCherry-H40I30 IVTT is inhibited inside and outside leads to 1.4+-0.2 fold SmN ratio and a SmL ratio of 1.1+-0.1 (also see d (ii, iii)) **d, (i)** The left plot illustrates the total red fluorescence emission of the 50 μl PMBC solution (fivefold diluted in buffer) after 5 hours mCherry-H40I30 IVTT inside PMBC (i), recorded with a mulitplate reader (see Supplementary Table 4). The red fluorescence emission (at 610 nm) of the PMBC solution is significantly different (p < 0.01) when compared to 10 mM Tris-HCL buffer solution (blue) and the control (no IVTT) where IVTT is inhibited outside and inside (red) **(ii)**. The middle plot illustrates the mean SmL ratio of line scans through ten PMBC for the three conditions IVTT outside and inside, IVTT inside and no IVTT (see Supplementary Table 3). The SmN ratio of IVTT inside is significantly different (p < 0.01) when compared to both other IVTT condition. **(iii)**The right plot illustrates the mean SmL ratio of line scans through ten PMBC for IVTT outside and inside, IVTT inside and no IVTT. The SmL ratio of IVTT inside is significantly different (p < 0.01) when compared to the both other IVTT conditions. Error bares in d represent the s.d. of 4 measurements of one replicate (i) and the s.d. of 10 independent vesicle line scans (ii) and (iii). The scale bars in b and c correspond to 5 μm. All assays where independently replicated at least twice.

## Conclusion

Taken together we present a protocell model composed of an amphiphilic protein membrane based on prebiotic aa that we propose as origin for the first self-sustaining entities preceding phospholipid-based compartmentalisation.

Considering the prevalent environmental conditions on the prebiotic earth, the stability of PMBCs and most notably their simple building block synthesis (compared to lipids) highlights their significance as promising protocell model. The compatibility and integration of prevalent protocell membrane constituents (phospholipids, fatty acids and fatty alcohols) to the investigated PMBC demonstrates the plausible transition of a prebiotic protein membrane based protocell towards current phospholipid membrane based cells. The encapsulation of functional proteins or whole PMBCs into one another illustrates the possibility of subcompartmentalization and spatial separation of reactions within one PMBC (Fig. 1).

We could show that prebiotic PMBC accommodate not only anabolic ligation reactions but most notably DNA encoded synthesis and subsequent incorporation of their membrane constituents as major step towards self-sustaining systems. Our findings support that prebiotic PMBC represent a new type of protocell model enabling to construct simple artificial cells and to reveal plausible structural and catalytic pathways to the emergence of lipid-based compartmentalization. Even though the amphiphilic peptides employed in this work are accessible under prebiotic conditions, the minimum length of amphiphilic proteins has to be investigated. Employing the minimum length of prebiotic amphiphilic peptides while preserving the three dimensional structure forming capacity will substantiate our hypothesis.

The simplicity and multifunctionality of our amphipathic building units are important prerequisites for their emergence as constituents of first prebiotic protocells. Rather more speculative, prebiotic amphipathic protein membranes might not only play a role as compartment boundary for self-replicating RNA molecules or early chemical reaction networks but they may contribute to the emergence of protocells in multiple ways: In extant cells the dynamic multifunctionality of proteins, notably the integral membrane proteins, is indispensable, implying a significant role within the prebiotic protocellular membrane. The plasma membrane of extant cells is composed of roughly equal parts of proteins and lipids whereas proteins are crucial for main tasks such as selective permeability, transport, recognition, regulation etc. One scenario could have been that lipids as boundary molecules inserted into PMBC as subsequent transition event. The next step for the PMBC protocell model would be the demonstration of prebiotic PMBC constituted of autocatalytic amphipathic peptides. This self-replicating activity of the compartment forming unit is crucial for the evolution of two different replicating systems: the informational genome (RNA replicase) and the compartment in which it resides. A self-replicating protocell model based on peptides is considerably more conceivable then lipid based systems. Dipeptides such as diglycines^54^, histidyl histidine^55^ Ser-His^56^ can effectively catalyze their own synthesis or catalyse peptide bond formation. In addition, autocatalytic synthesis of glycine to oligo glycine in a simulated submarine hydrothermal system^25^ emphasizes the mulitfunctionality of peptides and therefore their potential as self-replicating compartment constituents. Accordingly the proposed PMBC protocell model represents an important step towards a self-replicating protocell compartment. Furthermore, the association and coevolution of RNA precursor together with positively charged amphiphilic peptides e.g. K40I30 constituting one PMBC membrane is reasonable as Koga et al. have shown that poly(Lys) peptides associates with nucleotides to functional micro droplets^19^. In addition, Bergstrom et al. reported delivery of chemical reactivity to RNA and DNA by a specific short peptide (AAKK)4^57^.

In summary a dynamic protocell membrane constituted of simple prebiotic amphipathic peptide molecules possess major advantages when compared to a protocell enclosed by lipid membranes. First, the synthesis of peptide membrane constituents,^56,58^ is far more simple, second, the membrane itself can potentially exhibit autocatalytic and catalytic activity,^57,59^ and potentially self-replicate under simple conditions. Third, an associated coevolution of the informational genome (e.g. RNA), functional catalytical pathways (proteins) and the structural entity of the PMBC is conceivable. We believe that the presented PMBC protocell model enables new perspectives and provides new insights into the emergence of self-sustaining protocells on the pathway to living systems.

## Acknowledgement

The authors thank the BMBF for financial support and the Zentrum für Biosystemanalyse (ZBSA) for providing the research facility. We are grateful to the EM facility of the Department of Neuroanatomy Freiburg and thank Barbara Joch and Prof. Andreas Vlachos for help with TEM imaging. We thank the lab of Peter Kele for synthesizing and providing Combo Rhodamin. We thank the lab of Günter for providing the IVTT mix. We are grateful to P. G. Schultz, TSRI, La Jolla, California, USA for providing the plasmid pEVOL pAzF.

## Author Contributions

A.S., M.C.H. and S.M.S. conceived the project. A.S. and M.C.H. designed and performed the experiments. M.C.H. designed and cloned the used constructs. M.C.H. and A.S. conceptualized the publication, A.S. wrote the initial draft and the final publication together with M.C.H. S.M.S commented and discussed on the manuscript. A.S. and M.C.H contributed equally to the publication.

